# Computational modeling of cell signaling and mutations in pancreatic cancer

**DOI:** 10.1101/2021.06.08.447557

**Authors:** Cheryl A. Telmer, Khaled Sayed, Adam A. Butchy, Kara Bocan, Christof Kaltenmeier, Michael Lotze, Natasa Miskov-Zivanov

**Affiliations:** Carnegie Mellon University, Pittsburgh PA; University of Pittsburgh, Pittsburgh PA; University of Pittsburgh School of Medicine, Pittsburgh PA

## Abstract

Published research articles are rich sources of data when the knowledge is incorporated into models. Complex biological systems benefit from computational modeling’s ability to elucidate dynamics, explain data and address hypotheses. Modeling of pancreatic cancer could guide treatment of this devastating disease that has a known mutational profile disrupting signaling pathways but no reliable therapies. The approach described here is to utilize discrete modeling of the major signaling pathways, metabolism and the tumor microenvironment including macrophages. This modeling approach allows for abstraction in order to assemble large networks to capture numerous facets of the biological system under investigation. The Hallmarks of Cancer are represented as the processes of apoptosis, autophagy, cell cycle progression, inflammation, immune response, oxidative phosphorylation and proliferation. The model is initialized with pancreatic cancer receptors and mutations and simulated in time. The model portrays the hallmarks of cancer and suggests combinations of inhibitors as therapies.

## Introduction

Cancer as a genetic disease was suggested in 1902 by Theodor Boveri (Boveri 2008). Since that time there have been many therapies, many deaths and also many survivors. Although some cancers such as breast and colon can be managed, others such as glioma and pancreatic cancer have very poor survival rates. It is estimated that 44,330 people will die of pancreatic cancer in 2018 (Society 2019). Pancreatic cancer has early KRas activating mutations followed by TP53 and CDN2A inactivating mutations in the majority of tumors (Zeitouni, Pylayeva-Gupta et al. 2016, Raphael, Hruban et al. 2017). There are no available drugs to target KRas activation or restore tumor suppressor function and therefore survival of patients has not improved. Modeling is an approach that can improve understanding of a complex system, test numerous combinations of drugs and offer novel therapeutic options.

There are different types of models that can be utilized including statistical models that indicate correlations between tumor features and treatments (reviewed by (Altrock, Liu et al. 2015, Azuaje 2016), agent-based models (Norton, Wallace et al. 2017) and more detailed PDE (Simbawa 2017) and ODE models (Murphy, Jaafari et al. 2016). Each of these approaches has strengths and weaknesses related to available knowledge, size of the model, computational time and methods for analysis (Bartocci and Lió 2016). Cell signaling pathways are used by cells to transduce extracellular signals for sensing and adapting and are altered in disease states. Signaling can influence metabolism, cell survival, or cell death through complex combinations of protein degradation, post-translational modifications, protein translocations and gene expression. In cancer, the aberrant behavior of signaling pathways is observed however actual causes are difficult to determine due to the complexity of the networks of regulation that results from crosstalk, feedforward and feedback loops. Timing of these events and the outcomes of treatments vary so much that drug development trials often fail due to inaccurate predictions of the effects of inhibitor drugs. The approach described here is able to capture features and timing of communication of the tumor and microenvironment that are involved in tumor progression, including macrophages and pancreatic cells, extracellular matrix, and organelles within cells involved in signaling, gene regulation and metabolism. This representation of signaling pathways is then parameterized and simulated stochastically (Sayed, Kuo et al. 2017). Spatial and temporal components are included as subcellular organelles and translocations between organelles and cells (Sayed, Telmer et al. 2017). The influence of regulators is described as positive or negative depending on their effect on activity or amount of an element and elements are updated to new states using functions determined from the relationships of the regulators (Miskov-Zivanov, Turner et al. 2013). This approach uses discrete variables to describe the state of the element as low, medium or high. Discrete modeling methods include Boolean frameworks, logical models and Petri nets and have been applied to cancer signaling pathways (Chowdhury, Pradhan et al. 2013, Fumia and Martins 2013, Hu, Gu et al. 2015, Lu, Zeng et al. 2015, Cho, Park et al. 2016). The abstraction allows for more comprehensive collections of elements and inclusion of cellular processes, and this model is an extension of previous models of pancreatic cancer cells (Gong 2013, Wang, Miskov-Zivanov et al. 2016). The macrophage elements can be removed to create a pancreatic cell line model and a melanoma cell line would incorporate different mutations.

The cancer model demonstrates the development in time of the ground truth properties of cancer described in “The Hallmarks of Cancer” 2000 and 2011 (Hanahan and Weinberg 2000, Hanahan and Weinberg 2011). These are represented in the model as the cellular processes of apoptosis, autophagy, cell cycle progression, immune response, inflammation, oxidative phosphorylation and proliferation. Components of pancreatic cancer cell (PCC) pathways were obtained from papers cited in the Hallmarks papers (see Table 1). Tumor associated macrophages have been shown to be involved in the progression from pancreatitis to pancreatic cancer (reviewed by (Valilou, Keshavarz-Fathi et al. 2018)) and therefore were incorporated for simplified immune feedback in the tumor microenvironment.

**Table 1.** This table shows the numbers of references used to build the DySE model of pancreatic cancer from the Hallmarks of Cancer (excluding cancer stem cells, endothelial cells, pericytes, fibroblasts, stromal cells) and then the number of papers that were cited in those articles.

| Process | Paper ID | No. |
| --- | --- | --- |
| **Hallmarks of Cancer 2000** | PMID10647931 | 103 |
| **Hallmarks of Cancer 2011** | PMID21376230 | 237 |
| **Autophagy** | PMC2696814 | 101 |
| **Bcl-2 and apoptosis** | PMC2930981 | 144 |
| **DNA damage response** | PMC2988877 | 232 |
| **Glycolysis** | PMC2476215 | 61 |
| **HMGB1 signaling** | PMID20969478 | 272 |
| **Immune evasion** | PMC2891151 | 86 |
| **Immune inflammation** | PMC2865635 | 96 |
| **Inflammation** | PMC2866629 | 158 |
| **mTOR** | PMC3193604 | 27 |
| **Necroptosis** | PMID19109884 | 10 |
| **Oxidative stress** | PMC4204162 | 50 |
| **Ras family and exocyst** | PMID 12074877 | 24 |
| **Receptor Tyrosine Kinases** | PMC2914105 | 142 |
| **ROS signaling** | PMC3454471 | 149 |
| **Senescence** | PMC3672965 | 58 |
| **Telomerases** | PMC3003493 | 161 |
| **Warburg effect** | PMC2849637 | 47 |

This paper describes a computational model of cancer; a discrete model of signaling, metabolism and the pancreatic tumor microenvironment including macrophage cells. The model was assembled manually so that methods could be developed for computers to automatically assemble models from machine reading of scientific literature. The model organization and contents, modules within the model, and simulation results are followed by a discussion of the process, model applications and limitations. This directed, causative modeling framework incorporates knowledge and can perform reasoning which are both elements of artificial intelligence that are different from machine learning. The automation of the model creation and analysis will allow for a greater understanding of large complex systems.

## The Model

### Model elements

Major signaling molecules and pathways and their mechanistic, causal interactions were extracted from the Hallmarks of Cancer and citations within (see Table 1). At the plasma membrane are proteins that transduce extracellular signals to intracellular signals and transport molecules across the membrane. Extracellular ligands bind to and activate receptors and nutrients enter via transporters. The mitochondrial contribution to energy metabolism and reactive oxygen species generation are included. Proteins translocate between the cytoplasm and nucleus and RNA is generated prior to the presence of the proteins where the genes are the targets of transcription factors. A summary of the elements is shown in Table 2. The model spreadsheet is available <u>here</u>.

**Table 2.** Summary of the elements in the model.

| Process | Cellular feature | No. |
| --- | --- | --- |
| **Cell type** | Pancreas | 181 |
|  | Macrophage | 55 |
| **Location in the cell** | Nucleus | 105 |
|  | Cytoplasm | 60 |
|  | Plasma Membrane | 46 |
|  | Mitochondria | 4 |
|  | Endoplasmic reticulum | 1 |
| **Extracellular signaling** | Extracellular | 15 |
| **Other extracellular** | Extracellular | 5 |
| **Element type** | Protein | 49 |
|  | Protein family | 87 |
|  | Protein complex | 9 |
|  | mRNA | 33 |
|  | Gene | 34 |
|  | Chemical | 5 |
|  | Chemical family | 3 |
|  | Biological process | 15 |
| **Protein functions** | Receptors | 20 |
|  | Transcription factors | 16 |
| **Total elements** |  | 236 |

### Standardized notation for variable names in a tabular format

A tabular format is used for the entry of information about the elements (Sayed, Telmer et al. 2017). Proteins and compounds to be included in the model were assigned variable names and unique IDs, UniProt for proteins, HMDB for chemicals and GO for cellular locations and processes. As part of the abstraction the IDs are listed. Currently, only protein families and complexed are listed however future versions include all known interacting proteins that are included for a specific node. The variable names of elements are of the standard form ELEMENTtype_locCELLTYPE and begin with a trivial ELEMENT name that is followed by a designation of the type of biomolecule, chemical (che) or protein, and proteins are assigned (pn) when the active unit is a monomer or homodimer, protein family (pf) when there are multiple homologous gene products or protein complex (pc) when multiple different subunits form a complex. The category is then followed by an underscore (_) and the cellular location (loc) is entered as cyto (cytoplasm/cytosol), nuc (nucleus), PM (plasma membrane), mito (mitochondria), ER (endoplasmic reticulum) and ex (extracellular space). And finally cell type pancreatic cancer cell (PCC) or macrophage (MAC).

### Mutations

Mutations are added as element variables (MUTPROTEIN) and added as positive or negative regulators of the gene, RNA and protein matching their mutant activity.

### Positive and negative regulators

Signaling networks relay information using protein-protein interactions (PPIs) including binding events, translocations from one cellular compartment to another, protein synthesis, degradation and post-translational modifications (PTMs). PTMs include phosphorylation/dephosphorylation, acetylation/deacetylation, methylation/demethylation, ribosylation/deribosylation, hydroxylation, glycosylation, sumoylation/desumoylation, farnesylation, and ubiquitination/deubiquitination. These PPIs can activate or increase the activity or amount of target proteins or they can inhibit or decrease the activity or amount of target proteins. The sign of the interaction represents how the regulator influences the regulated element eg. if the phosphorylation of Protein B by Protein A inhibits the activity then Protein A is a negative regulator of Protein B.

### Regulator notation

In the tabular format are columns for positive and negative regulators of the variables/elements so that if Protein A phosphorylates and activates Protein B then Protein A is a positive regulator of Protein B. Additionally, if there are multiple regulators they are entered in a notation to represent OR, AND, and enhancing pair (Sayed, Telmer et al. 2017). If there are multiple regulators such that Protein A or Protein B will positively regulate, then they are comma separated, A,B, where an AND relationship has the regulators in parentheses (A,B). The enhancing pair notation is used where a second element enhances the activation of an activator. This notation is often used to have the RNA of protein B increasing the activation of A on B and is written {A}[Brna] as positive regulators of protein B and shows that by increasing the amount of B we expect the activation to increase but only if A is present ie. A is necessary.

### Regulator motifs

This notation is used to describe the relationships between the regulators and their targets and many show up in motifs as for receptor activation when ligand AND receptor are necessary for the active form of the receptor ((ligand, receptor) are positive regulators of receptor_ACT). A translocation motif has the cytoplasmic version of the protein being activated and entering the nucleus to serve as a transcription factor that then activates a gene regulatory motif (Sayed, Telmer et al. 2016). This motif has the gene activated and then forming an RNA and then the protein. These standardized motifs guide the user and serve to approximate the timing of signaling events. In the model, signals are transduced through 20 receptor motifs, 10 translocation motifs (ERK, HMGB1, MDM2, NFkB, NRF2, p38MAPK, ROS, SMAD3, STAT1, STAT3) and gene expression motifs include 33 RNA species, 23 in the PCC and 10 in the MAC.

### Intracellular compartments and organelles

Cellular compartments allow for the specialization and regulation of cellular processes and transfer between compartments will influence the timing of signal transmission. Translocation motifs are implemented for proteins moving between compartments. These unique protein forms are used for instances where signal transduction affects only proteins at a specific location such as p53 in the cytoplasm is activated by NFkB in the cytoplasm and then goes on to inhibit AMPK (Tasdemir, Maiuri et al. 2008).

### Cellular processes

Cancer is characterized by alterations in cellular processes resulting from changes in signaling and gene expression. These processes reflect the ground truths as described by Hanahan and Weinberg 2000, 2011 (Hanahan and Weinberg 2000, Hanahan and Weinberg 2011). In the model the processes are influenced by inputs from signaling pathways and show the influences of these multiple pathways over time. The processes used as “ground truths” or “properties” that the model was constructed to analyze are apoptosis, autophagy, cell cycle progression, immune response, inflammation, oxidative phosphorylation (OXPHOS) and proliferation. Normally, cells have high CCA and OXPHOS and the other processes are low, however, cancer is characterized by low apoptosis, immune response and OXPHOS while other processes such as autophagy, cell cycle progression and inflammation are increased as well as proliferation which is the strongest indicator of cancer.

### Translation to logic rules

The table is then used to write logic rules (Sayed, Kuo et al. 2017).

### Initialization

The initial values of variables/elements are set to 0, 1 or 2 to represent Low, Medium or High level of activity/amount and are summarized in the spreadsheet of the model elements. All receptors are initialized at 1. In order to model the development of cancer, mutations are required, an activating RAS mutation at step 1000 and later at step 3000, inactivating mutations of p53 and CDN2A.

For cell line models the macrophage elements and regulators are omitted. Different tumors and cell lines harbor different mutations and therefore are changed to reflect the tumor origin when the information is available.

### Simulations

Logic rules are then used for multiple runs of a designated number of steps for the simulations and plots are created. Regulators are subject to logic rules for updating variables (Sayed, Kuo et al. 2017).

Traces were generated from 500 runs for 10,000 time steps.

## Modules

Cancer is a genetic disease and this is accounted for in the model by incorporating mutations and their effect on the protein either as activating or inhibiting. Represented in the model are several modules involved in cancer, including the major intracellular signaling pathways, metabolism, and the extracellular microenvironment with macrophages.

### Genotype

Pancreatic cancer genetics are not as diverse as many other forms of cancer. During cancer development there is an early activating KRas mutation followed by high rates of inactivating TP53 and CDN2A mutations these are incorporated into the model structure.

### Major signaling pathways in cancer

In order to gain understanding of cancer, the major signaling pathways and crosstalk are represented. Within cells there are often opposing or redundant effects so that a stimulus or ligand will activate one pathway to cause an effect, and another pathway to limit the activation. There has been an attempt to select and balance the effects included in the model. The interactions over time are often difficult to predict manually due to the complexity of the network however, machines can compute the outcomes in minutes. Each of the major signaling pathways is described below.

### RAS/ERK

Alterations in the Ras pathway are involved in numerous cancers including pancreatic cancer. The canonical activation is EGF binds to EGFR to activate it, then the receptor activates RAS which activates RAF which activates MEK which activates ERK and ERK translocates to the nucleus where it stimulates gene expression that leads to proliferation. In the model, the epithelial cell RAS is activated by EGFR, VEGFR, RAGE, TGFbR, SRC and IRS.

### TNFa/NFkB

NFkB in the cytoplasm is activated by the TNFa receptor, TNFAR, and also IL1R, PKCA and AKT. The NFkB then translocates to the nucleus to influence gene expression. In the model we abstract the NFkB pathway, such that the TNFa receptor activates TRAF2 which activates NFkB which is translocated. This representation simplifies TRAF2 phosphorylation and activations of IKK which then phosphorylates IkB (complex of IKKa,b,g that is complexed with NFkB complex (dimers of RelA/p65, RelB, c-Rel, p50 or p52)to keep it inactive), the phosphorylation of IkB promotes ubiquitination and targeting to the proteasome for degradation resulting in liberation of NFkB and allowing for translocation to the nucleus where it assembles with coactivators and RNA polymerase to induce gene expression. This pathway is also represented in the macrophage. There are many variations in the activators, the dimers in the NFkB complex and the coactivators that can be specified if one is interested in capturing more detailed knowledge of this pathway or comparing canonical and noncanonical signaling pathways.

### JAK/STAT

When interferons and interleukins bind their receptors they recruit a Janus kinase (JAK1, JAK2, JAK3 or TYK2) that then recruits and phosphorylates a STAT protein (STAT1, 2, 3, 4, 5A, 5B or 6). STATs then homo or heterodimerize, migrate to the nuclear pore and bind importins and are translocated into the nucleus where they act as transcription factors. STATs also recruit coactivators and undergo other PTMs that affect activity. The model has IL1, IL6, and interferon gamma that activate JAK/STAT signaling in the pancreatic cell and macrophage.

### TGFb/SMAD

When the TGFb receptor binds TGFb it recruits additional receptors to the complex and then phosphorylates SMAD2 and SMAD3, if the receptors bind bone morphogenic protein, BMP, the receptors phosphorylate SMAD1, SMAD5 and SMAD8. These phosphorylated SMADs then recruit SMAD4 to form a complex that is translocated to the nucleus to function as a transcription factor, expressing genes to decrease proliferation. The model currently has only the TGFb pathway.

### AMPK/mTORC1

The mTORC1 protein complex is involved in controlling protein synthesis in response to nutrient and redox conditions. The AMP activated protein kinase, AMPK, responds to energy status within cells. Low energy supply activates AMPK to phosphorylate TSC2, an inhibitor of mTORC1. The mTORC1 is also influenced by other signaling pathways including AKT. The mTORC1 inhibits autophagy and is mediated by several other protein complexes that are not included in the model.

### PI3K/AKT

The PI3K is also activated by receptor tyrosine kinases and initiates the conversion of PIP2 to PIP3 (dephosphorylated by PTEN) that then activates PDPK1 to phosphorylate AKT. The effects of AKT are numerous and in the model are activation of MDM2, mTORC1 and NFkB and inhibition of TSC2.

### ASK/JNK

It is important for the cell to control the redox state of the cytoplasm and ASK (apoptosis signaling kinase) is one of the kinases that responds to oxidative stress and TRAFs to then phosphorylate MEK 4/7 which phosphorylates JNK or MEK 3/6 and then p38MAPK (but not ERK) (The roles of ASK family proteins in stress responses and diseases (Hattori K 2009)).

### Metabolism

Integrally involved in cell function is the regulation of cell metabolism and while many metabolic processes utilize rapid allosteric controls we can incorporate components and reflect general processes. In the model, glucose and the glucose transporter are included, mitochondrial energy generating processes and lactate production are represented. Reactive oxygen species are important signaling molecules that result from inefficient electron transport that is observed in cancer.

### Glucose metabolism and lactate production

Modeling metabolism is different than modeling signaling cascades, the knowledge is more about amounts, and compounds are consumed and sent in one direction or the other. In this model lactate dehydrogenase is regulated by mitochondrial damage.

### ROS signaling and DNA damage response

The relationship between ROS and oxidative phosphorylation is complex, on the one hand ROS is produced when electron transport through the chain is compromised, and on the other hand ROS signals to decrease oxidative phosphorylation. In this model, ROS are generated from macrophages, and NADPH oxidase in pancreatic cells. If there is ROS in the cytoplasm it will cause mitochondrial damage.

### Extracellular environment and macrophages

Outside of the pancreatic cell, the extracellular environment, includes the extracellular matrix of protein and carbohydrate, lipid enclosed exosomes, growth factor and cytokine proteins, hormones, small molecules and ions, nutrients and metabolic byproducts which also influence physical features such as pH. The extracellular environment has components of blood, secretions from the pancreatic and immune cells such as macrophages and is the medium through which the cells communicate with each other. In the model is a simplified version of a macrophage including signaling pathways within the cell. An additional module within the model is the activation of exosome transfer to macrophages by RAS and the transfer of RAS protein to the macrophages.

### Macrophage

These immune cells are resident in tissues and are involved in inflammation and tissue repair. The macrophage in the model is quite simple with only 55 elements including 11 receptors for CCL2, CSF1, interferon gamma, IL1, IL6, IL10, PL1L1, RAGE, TGFb, TLR4, and TNFa, and 11 genes for CCL2, interferon gamma and interferon gamma receptor, IL1B, IL1 receptor, IL6, IL6 receptor, IL10, STAT1, TGFb, and TNFa. Signaling utilizes JAK/STAT, SMAD, NFkB and ERK pathways. Proinflammatory and anti-inflammatory cytokines are released and compose the communications between cells.

### Ground truths

Analysis of the simulations was assessed by considering the trajectories of the biological processes extracted from the Hallmarks of Cancer, apoptosis, autophagy, cell cycle progression, immune response, inflammation, oxidative phosphorylation and proliferation.

### Simulation Results

This model of the development of cancer includes representations of activation of signaling pathways, interactions with the tumor microenvironment, the effect of mutations and inhibitors. The model has cancer is dependent upon mutation, injury shows moderate increases in proliferation but not high proliferation. When the simulation begins there are no mutations and the initial values of the signaling pathways are as described above. Each of the processes behaves differently with injury showing intermediate effects. There is a small initial apoptosis, autophagy and immune response, cell cycle progression and inflammation increase slightly and OXPHOX decreases with proliferation increasing slightly. If however there is a mutation where an activating Ras mutation at 1000 steps, is followed by inactivating p53 and CDN2A mutations at 3000 steps, the ground truth properties of cancer are observed such that apoptosis and immune response remain low, autophagy, OXPHOS decrease and cell cycle progression, inflammation and proliferation increase (Figure 1).

**Figure 1.**
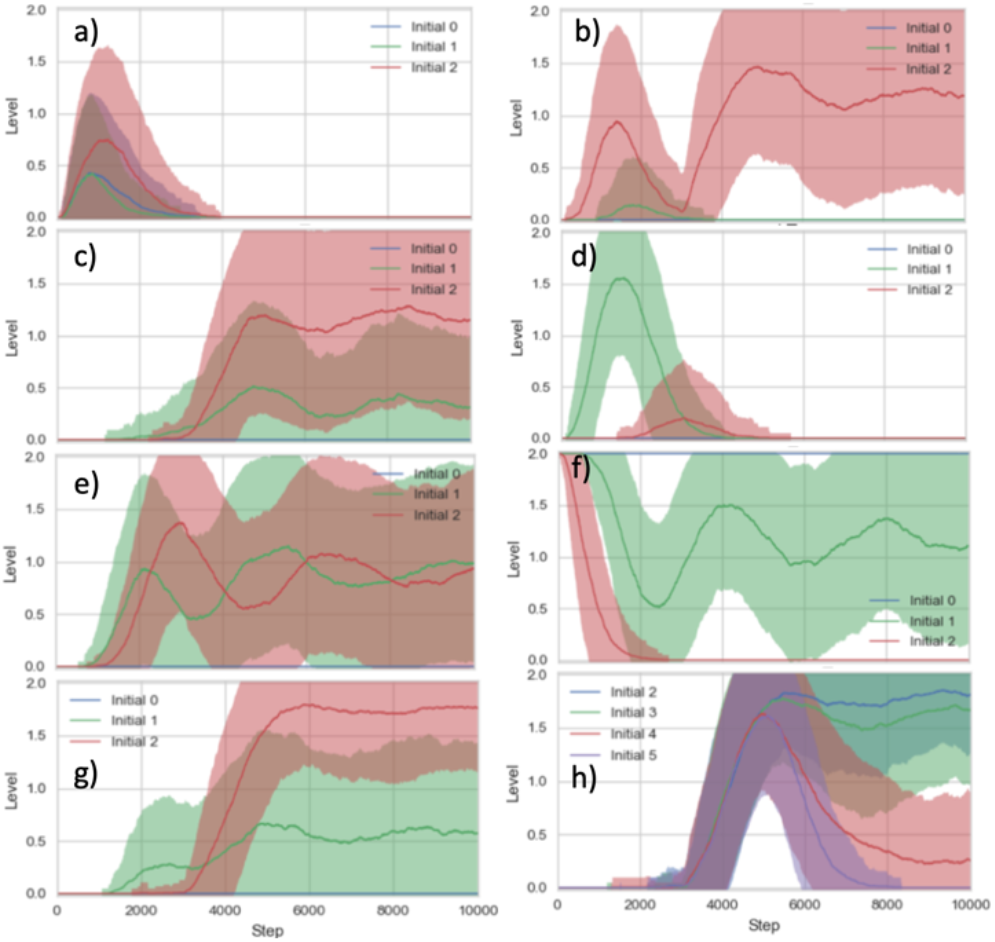
Plots of the activity level of processes a. apoptosis b. autophagy c. cell cycle progression d. immune response e. inflammation f. oxidative phosphorylation g. proliferation h. proliferation with inhibitors. The scenarios are shown as blue, Initial 0 is normal, green, Initial 1 is with injury and red, Initial 2 is with KRas, TP53 and CDN2A mutation, Initial 3 is with MEK inhibitor, Initial 4 is with PI3K inhibitor and Initial 5 is with MEK and PI3K inhibitors.

### Intervention scenarios, MEK and PI3K inhibitors

The traces in Figure 1h show the results for proliferation with inhibitors added. MEK inhibitor alone only decreases proliferation slightly. PI3K inhibitor results in a larger decrease and the addition of both decreases proliferation completely.

### Cancer Cell Lines

Many research studies employ cell lines to screen compound libraries or investigate the effects of small molecule treatments. In order to have a model for simulating cell lines we remove the macrophage elements and regulators. For related cell lines the mutations corresponding to the cancer cell lines are modified. Models of pancreatic cancer and melanoma cell lines have been developed.

## Discussion

### Manual assembly of a complex model

A model is a standardized representation of knowledge about a subject and can be at many different levels of abstraction and detail. Here the system is cancer, the type is pancreatic cancer, the level is cellular signaling and the approach is based on circuit design and logic and utilizes rules for updating node states that are a function of the regulators. The signaling pathways are represented by key molecules at nodes that then function to regulate other nodes positively or negatively. The system is simulated repeatedly for a number of time steps and average dynamic behavior of the elements is produced. For many of the interactions there are decisions about the level of abstraction.

### Abstraction

The tumor microenvironment is also an extremely large complex network that includes different cell types, cell signaling pathways, cell metabolism and involves multiple cell compartments. In order to account for interactions between these numerous features where there is often incomplete information about rates, this modeling framework utilizes abstract representations at the nodes and discrete levels of low, medium and high. With this scaffold, modules of elements and paths can be incorporated to expand the model for answering specific detailed questions.

Dimerization (Amoutzias, Robertson et al. 2008), homo and hetero, accessory proteins, chaperones, and other protein-protein interactions and modifications are all strategies used by cells to regulate signaling networks. In this model these mechanisms are abstracted at the nodes such that the influence of one element on another is represented in the update rule for the downstream regulated element. This model was built manually and therefore human judgement was used to decide how much detail to incorporate. In order to automate this procedure it is necessary to have automatic methods for adding and evaluating the addition of new information.

Current research is producing more and more detailed experimental results. For example, many years and numerous studies have shown that EGF binds to EGFR, the receptor dimerizes, autophosphorylates and recruits Shc and Grb2 which then recruit SOS from the cytosol and the complex activates Ras which in turn activates Raf. Raf then phosphorylates MEK and MEK phosphorylates ERK (Lake, Corrêa et al. 2016). There are then two major feedback loops, one post-translational and one transcriptional. The model described here does not include adapter proteins of signaling cascades, instead highlights major components, understanding that this is not a complete representation but a simplified version, an abstraction of the cascades. For detailed investigations of specific signaling pathways more components can be added to represent deeper mechanistic feedbacks and understanding while maintaining the ground truths of cancer. As models get larger the computational time required for the simulations also increases.

The cancer model represents the development of cancer over years and the cell lines show experimental simulations representing days. As methods improve these features will become more defined and standardized. Combined with the vast amounts of biological data that are being generated, the biomedical field can benefit from modeling to provide a scaffold for this knowledge and simulation of the dynamics is important for generating and testing hypotheses and making predictions and improving our understanding of these complex systems.

### Baseline model

Proteins and their regulatory relationships were interpreted from “The Hallmarks of Cancer” (Hanahan and Weinberg 2000, Hanahan and Weinberg 2011) and adapted to pancreatic cancer. The pancreatic cancer cell network was then extended with elements of ROS signaling pathways, exosomes, HMGB1, and a minimal abstract representation of macrophages. Extensions to the model were incorporated as paths so that additions were regulated by an element in the model at the start of a path and the element at the end of the path was a regulator of an element in the model. If genes were involved then RNA and protein forms were also included. To create these paths, iterative searches of the literature using different adjectives or processes were used to find information about how a protein was connected to the model. Computers can read much larger volumes of literature and therefore this can be automated by searching the elements of interest, collecting the papers mentioning the elements, adding in extracted relationships and measuring the effect on the model outcomes.

This model represents genotype, multiple signaling pathways, receptors, organelles, translocations and biological processes. It can serve as a baseline to consider specific pathways in detail while still accounting for interactions with other cell signaling pathways. While not comprehensive, we have attempted to capture the influential molecules of the major signaling pathways and some of the crosstalk and feedback and feedforward loops. There are instances where the data is currently conflicted such as SMAD4 mutation in pancreatic cancer (Di Marco, Astolfi et al. 2015) and therefore was not included here and will be the subject of future studies. As knowledge increases it can be added to this model in order to represent cancer more accurately and can be expanded upon for answering specific questions. This baseline is easily modified to represent cell lines, a common experimental model for testing of therapeutic compounds.

Simulations of tumors and the microenvironment coupled with data from the tumors will enable precise and personalized tumor treatment in order to improve quality of life and survival for years beyond a cancer diagnosis. Artificial Intelligence is not really artificial, it is the automation of learning and reasoning. Synthetic Biology is synthetic in the “synthesis” use of the word and using intelligent computational modeling to aid in the understanding of large complex systems will result in advances in not only in biology and medicine, but also in environmental sciences, agriculture and economics.

## Acknowledgements

We would like to than our collaborators Peter Sprites for many insightful comments. This work is supported by DARPA grants W911NF-17-1-0135 and W911NF-18-1-0017-(72314-NS-DRP) awarded to N. Miskov-Zivanov.

